## Supplementary figures and images for "Axonal distribution of mitochondria maintains neuronal autophagy during aging via eIF2β"

### Figure S1

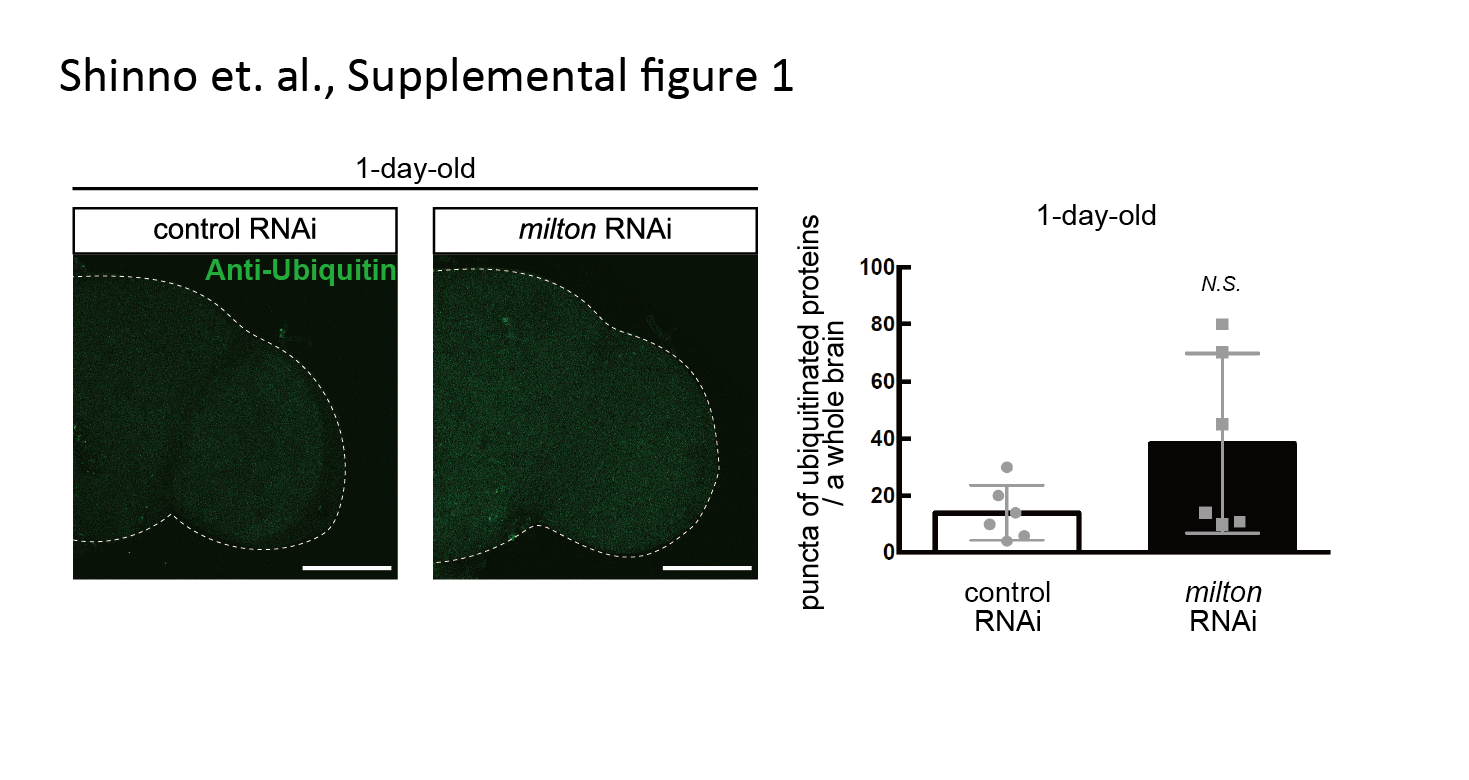

### Figure S2

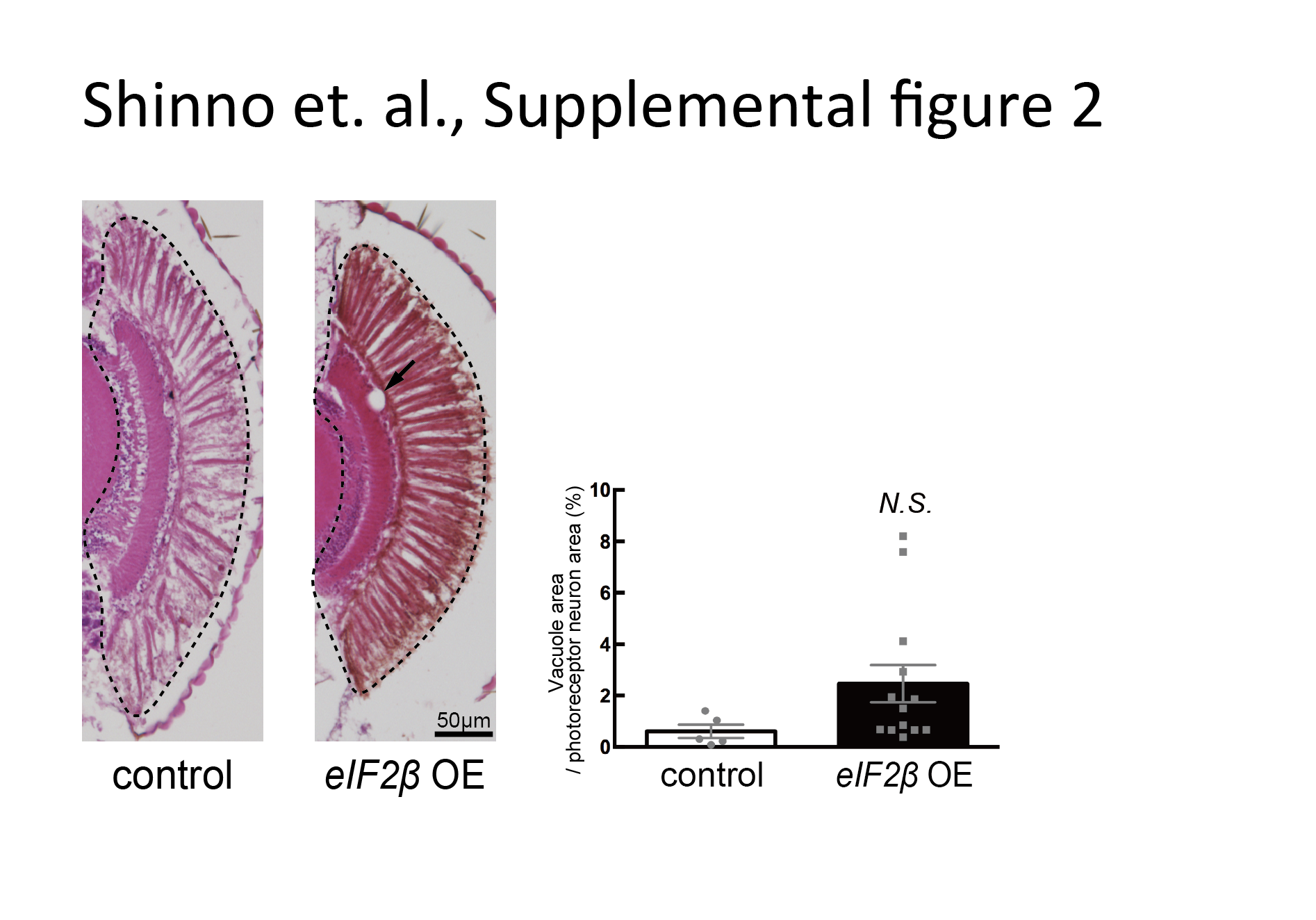
